# A genome-wide association study finds novel genetic associations with broadly-defined headache in UK Biobank (N = 223,773)

**DOI:** 10.1101/217786

**Authors:** Weihua Meng, Mark J Adams, Harry L Hebert, Ian J Deary, Andrew M McIntosh, Blair H Smith

**Affiliations:** Division of Population Health Sciences, School of Medicine, University of Dundee, Dundee, UK, DD2 4BF; Division of Psychiatry, Edinburgh Medical School, University of Edinburgh, Edinburgh, UK, EH10 5HF; Centre for Cognitive Ageing and Cognitive Epidemiology, Department of Psychology, University of Edinburgh, Edinburgh, UK, EH8 9JZ

**Keywords:** headache, genome-wide association study, *LRP1*, UK Biobank, tissue expression

## Abstract

Headache is the most common neurological symptom and a leading cause of years lived with disability. We sought to identify the genetic variants associated with a broadly-defined headache phenotype in 223,773 subjects from the UK Biobank cohort. We defined headache based on a specific question answered by the UK Biobank participants. We performed a genome-wide association study of headache as a single entity, using 74,461 cases and 149,312 controls. We identified 3,343 SNPs which reached the genome-wide significance level of *P* < 5 × 10^−8^. The SNPs were located in 28 loci, with the top SNP of rs11172113 in the *LRP1* gene having a *P* value of 4.92 × 10^−47^. Of the 28 loci, 14 have previously been associated with migraine. Among 14 new loci, rs77804065 with a *P* value of 5.87 × 10^−15^ in the *LINC02210-CRHR1* gene was the top SNP.

Positive relationships (*P* < 0.001) between multiple brain tissues and genetic associations were identified through tissue expression analysis, whereas no vascular related tissues showed significant relationships. We identified several significant positive genetic correlations between headache and other psychological traits including neuroticism, depressive symptoms, insomnia, and major depressive disorder.

Our results suggest that brain function is closely related to broadly-defined headache. In addition, we also found that many psychological traits have genetic correlations with headache.

## Introduction

Headache is the most common neurological symptom, with a life time prevalence of over 90% in the general population in the UK (1). It represents 4.4% of consultations in primary care and 30% of outpatient consultations in neurology (2,3).

According to the International Headache Society, headache can be generally divided into two categories: primary headache, if not associated with another disorder; and secondary headache if associated with an underlying medical illness (4). Primary headaches mainly include migraine, tension-type, and cluster headaches. Secondary headaches include any head pain caused by infection, neoplasm, head injury, some metabolic disorders, or drugs (4).

In a comprehensive review of population-based epidemiological studies of headache, the global prevalence of recurrent headache in all ages was found to be 46% for all headaches, including 11% for migraine and 42% for tension-type headache (5). Tension-type headache is the most prevalent type of headache, whereas migraine is the most disabling (6).

Migraine affects around 6 million people in England in the age range 16-65 and it is the sixth cause in terms of years of life lost to disability according to the Global Burden of Diseases 2013 (7). Migraine costs the National Health Service (NHS) almost £2 billion per year (8). It presents with recurrent headache attacks and/or hypersensitivity to light and sound. Around one third of migraineurs experience an aura, which are transient neurological symptoms mostly involving the visual system (9).

Family studies and twin studies have suggested that both migraine and tension-type headache are heritable traits with a heritability over 40% (10,11). Recently, genome-wide association studies (GWAS) have identified many genetic loci associated with migraine (12–16). A GWAS meta-analysis of 375,000 patients involving 22 centres has identified 38 genetic susceptibility loci for migraine with the *LRP1* region in chromosome 12 being the most strongly associated (17). Along with other GWAS on migraine, the total number of loci identified to be associated with migraine is currently 47 (18). No GWAS have been performed for tension-type headache so far.

There are several phenotypic associations between headache and metabolic, psychological, and other factors such as obesity (19,20). Genome-wide association studies provide a potential route to discover genetic correlations with other complex traits and diseases that in turn may provide clues to shared genetic architectures and aetiologies (21).

To identify the genetic variants associated with headache, we conducted this GWAS using the UK Biobank cohort which has never been contributed to genetic studies of headache including migraine. We used a broadly-defined headache phenotype, the one available in the UK Biobank dataset. Secondly, we sought to test for shared genetic associations with other complex traits and diseases using linkage-disequilibrium score regression (22).

## Results

### GWAS results

During the initial assessment visit (2006-2010), at which participants were recruited and consent was given by 501,708 UK Biobank participants, the specific pain question (see the materials and methods section for details) received 775,252 responses to all options. Among these responses, the number of participants who selected the ‘Headache’ option was 102,994 (cases), and the number of participants who selected the ‘None of the above’ option was 197,149 (controls). We further removed those whose ancestry was not white British (n = 22,694) based on principal component analysis, those who were related to one or more others in the cohort (a cut-off value of 0.025 in the generation of the genetic relationship matrix) (n = 52,166), those who were also in a Psychiatric Genomics Consortium MDD cohort (n = 597), and those who failed QC (n = 913). Thus we finally identified 74,461 cases (27,350 males and 47,111 females) and 149,312 controls (71,480 males and 77,832 females) for the GWAS association analysis. After quality control, there were 9,304,965 single nucleotide polymorphisms (SNPs) for the GWAS analysis.

The clinical characteristics of these cases and controls are summarised in Table 1. There were statistical differences (*P*<0.001) in age, sex and body mass index (BMI) between cases and controls.

**Table 1.** Clinical characteristics of headache cases (74,461) and controls (149,312)

| Covariates | Cases | Controls | *P* |
| --- | --- | --- | --- |
| Sex (male:female) | 27,350: 47,111 | 71,480: 77,832 | <0.001 |
| Age (years) | 54.38 (7.95) | 56.93 (7.97) | <0.001 |
| BMI (kg/m<sup>2</sup>) | 27.50 (5.05) | 26.66 (4.30) | <0.001 |

BMI: body mass index
A chi-square test was used to test the difference of gender frequency between cases and controls and an independent t test was used for other covariates.
Continuous covariates were presented as mean (standard deviation).

We identified 3,343 SNPs which reached GWAS significance of *P* < 5 × 10^−8^ (Fig 1, Supplementary Table S1). These SNPs represented 28 independent loci including 14 newly-identified loci (Table 2).

**Figure 1.**
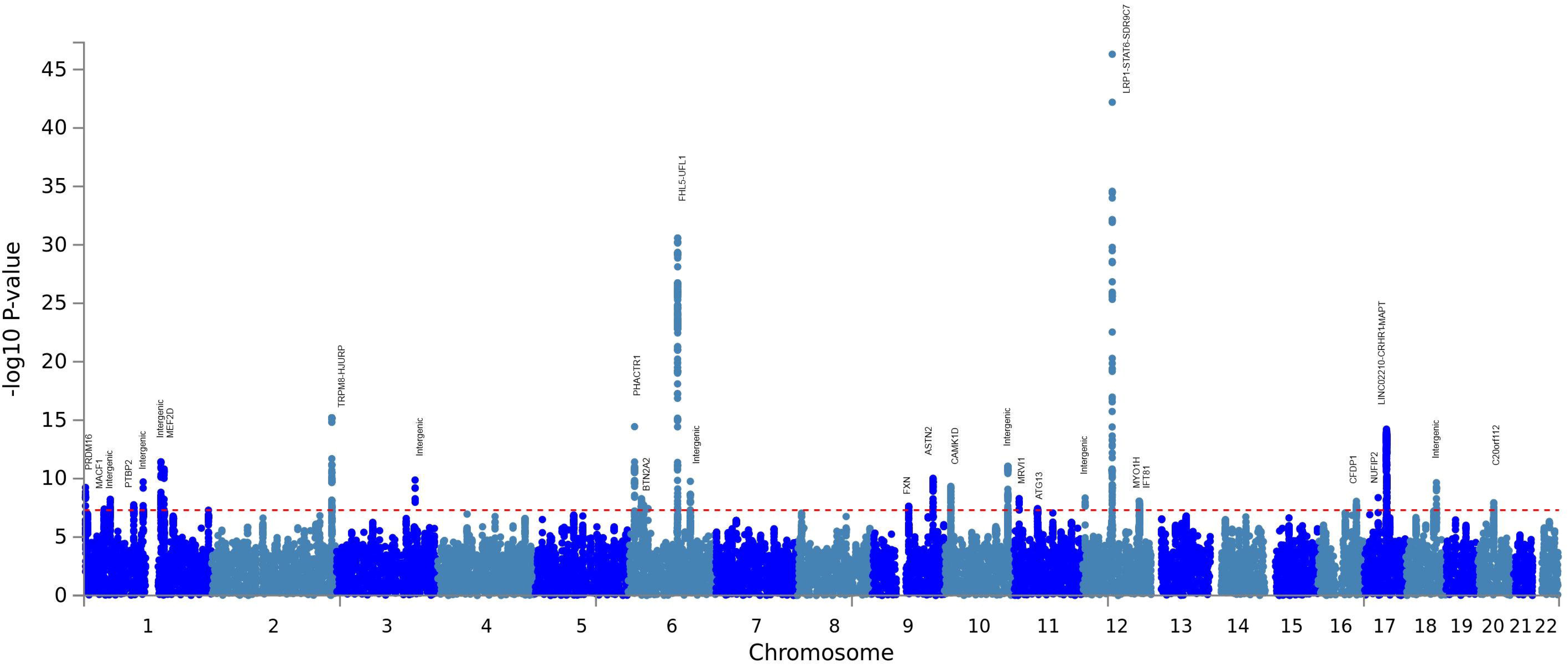
the Manhattan plot of the GWAS on headache using the UK Biobank cohort

**Table 2.**
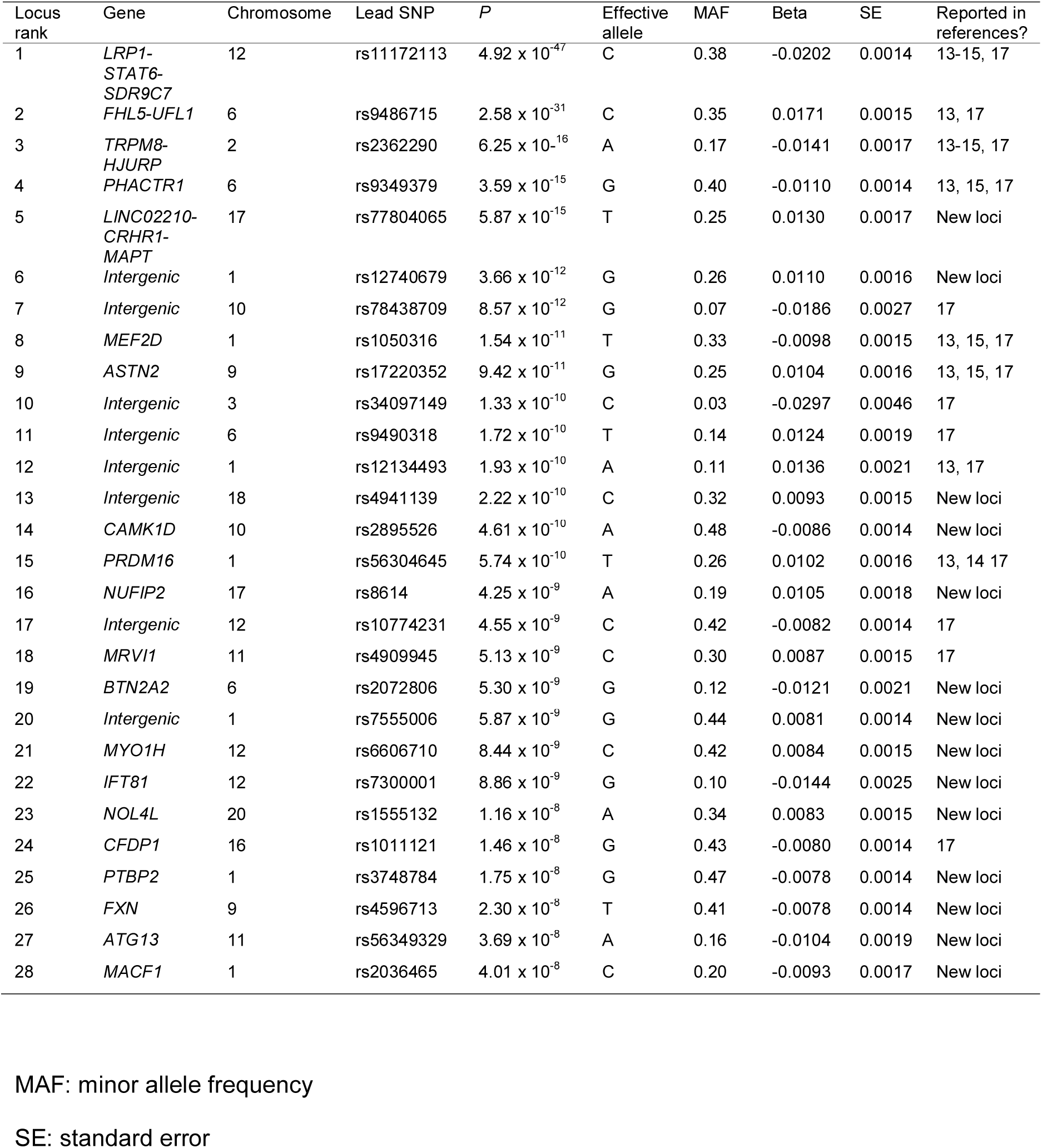
Summary of the 28 loci associated with broadly defined headache

The first cluster was in the LDL receptor related protein 1 (*LRP1*) gene in the chromosome 12 area with a lowest *P* value of 4.92 × 10^−47^ for rs11172113 (C allele, odds ratio (OR): 0.98). The second cluster was in the four and a half LIM domains 5 (*FHL5*) gene in chromosome 6 with a lowest *P* value of 2.58 × 10^−31^ for rs9486715 (C allele, OR: 1.02). Among the 28 loci, 14 of them were newly identified. Rs77804065 in the LINC02210-CRHR1 readthrough (*LINC02210-CRHR1*) gene with a *P* value of 5.87 × 10^−15^ (T allele, OR: 1.01) was the most strongly associated among the newly-identified loci. The Q-Q plot of the GWAS is shown in Supplementary Fig S1.

### Gene analysis, gene-set analysis and tissue expression analysis by FUMA

In the gene analysis by MAGMA integrated in FUMA, all the SNPs are mapped to 19,436 protein coding genes if the SNPs are located within genes. The default SNP-wise (mean) model for the gene analysis was applied. The signal transducer and activator of transcription 6 (*STAT6*) gene reached the lowest *P* value of 1.1 × 10^−46^, followed by UFM1 specific ligase 1 (*UFL1*) (*P* = 2.53 × 10^−26^), *FHL5* (*P* = 8.64 x10^−25^) and *LRP1* (*P* = 3.85 × 10^−23^). All genes (N = 160) with a *P* less than 3 × 10^−6^ (0.05/19436) are included in the Supplementary Table S2.

In the gene set analysis by MAGMA integrated in FUMA, a total of 10,894 gene sets were tested and a default competitive test model was applied. Gene sets of positive regulation of gene expression, positive regulation of transcription from RNA polymerase ii promoter, neurogenesis, and excitatory synapse reached a *P* value less than 0.0001, but not statistical significant of *P* < 5 × 10^−6^ (0.05/10,894). The top 10 gene sets from the analysis were included in Supplementary Table S3.

In the tissue expression analysis by GTEx integrated in the FUMA, average gene-expression per tissue type was used as gene covariate to test positive relationship between gene expression in a specific tissue type and genetic associations. Two types of tissue analysis were performed. One used 30 general tissue types and the other used 53 specific tissue types. In the expression analysis of 30 general tissue types, the tissue in the brain showed the most significant *P* value (*P* = 4.12 × 10^−6^), followed by the tissue from blood vessel (*P* = 0.014) (Supplementary Table S4). In the 53 specific tissue types, tissues from the brain cortex reached a lowest *P* value of 1.44 × 10^−5^ and most of the brain specific tissues also reached a significant *P* value of 0.001 (0.05/53). None of the vascular-related tissues reached significance. See Fig 2 and Supplementary Table S5.

**Figure 2.**
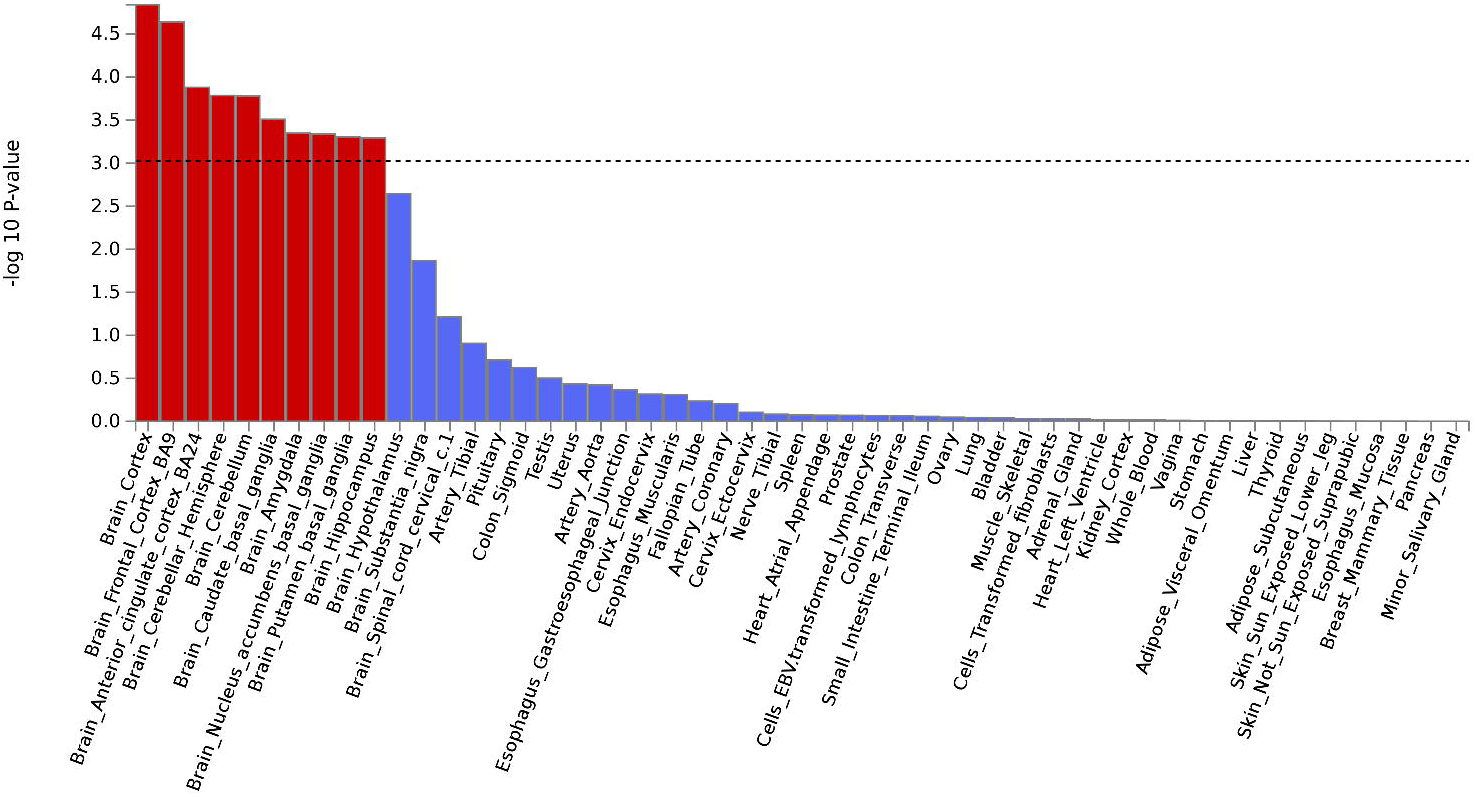
the tissue expression results on 53 specific tissue types by GTEx in FUMA The line indicates statistical significance of *P* = 0.001 (0.05/53)

### Genetic correlation analysis by LD hub

Through the genetic correlation analysis (see the materials and methods section for details), we identified multiple significant positive correlations for headache (Supplementary Table S6). The significant genetic correlations (rg) surviving multiple testing correction (*P* < 0.05/234) were: neuroticism (rg = 0.50, *P* = 2.24 × 10^−72^), depressive symptoms (rg = 0.52, *P* = 1.60 × 10^−46^), years of education (rg = −0.28, *P* = 5.25 × 10^−37^), maternal age at first delivery (rg = −0.32, *P* = 3.97 × 10^−29^), subjective wellbeing (rg = −0.37, *P* = 9.51 × 10^−19^), insomnia (rg = 0.42, *P* = 2.54 × 10^−18^), and major depressive disorder (rg = 0.39, *P* = 1.57 × 10^−11^).

## Discussion

We have performed a GWAS on broadly-defined headache as a single entity using the UK Biobank resource and found that variants in 28 loci were associated with having experienced headache within the last month to the extent that it interfered with usual activities. Evidence from tissue expression analysis showed that brain function is closely related to this broadly-defined headache. In addition, we found that headache was genetically correlated with a number of psychological factors, including those linked to a higher tendency toward experiencing negative emotional states, and shorter duration of education.

In this study, we defined headache cases and controls based on the responses by UK Biobank participants to a specific pain question. This question focused on headache occurrence, sufficient to cause interference with activities, during the previous month. We can therefore only treat headache as a global condition to perform GWAS. We can hypothesise, based on the reported population prevalence of each subtype (5,6), that tension-type headache will be the most common diagnosis among cases, followed by migraine, and that many will have experienced more than one type of headache. Whether the SNPs identified are associated with one or more of these diagnoses specifically, or with headache globally remains unclear until further research is conducted. It is important to note that half of the 28 loci were previously reported loci for migraine (Table 2).

The UK Biobank genetic resource is especially useful as a screening tool to test whether heterogeneous phenotypes such as headache have genetic components at all, as the UK Biobank has collected many heterogeneous phenotypes which need further genetic investigation. An example of such a GWAS approach was adopted by Deary et al. on self-reported tiredness (a heterogeneous phenotype) using the UK Biobank cohort (23). GWAS like this will help with genetic stratification analysis for heterogeneous phenotypes. This genetic discovery phase could be part of the partitioning or classification of broadly-reported or broadly-defined headache, improving our understanding of its aetiology, diagnosis, prognosis and the development of treatments.

The findings of this research will facilitate the next essential phases of research into the genetics and causes of headache. They will enable investigators to conduct adequate power calculations for future studies and will provide candidate loci to examine in replication studies. Even with our very heterogeneous group of cases, including many different headache subtypes, we were still able to identify numerous GWAS associations. Therefore, it can be expected that in a GWAS of a more intensively-phenotyped headache cohort or a subtype of headache such as tension-type headache for which no GWAS has yet been performed, genetic contributions could be more distinctive and easier to identify than in this study (for example, GWAS hits can be identified using fewer samples). Since our results are of GWAS significance, the genes (such as *LRP1* and *FHL5*) identified for migraine by other GWAS are also promising candidate genes for tension-type headache. The majority of the patients with migraine (over 90%) also suffer from tension-type headache (24), and it is possible that migraine and tension-type headache might share common genetic components.

In this GWAS, we have identified 28 loci for headache. One was in the *LRP1* gene with a lowest *P* value of 4.92 × 10^−47^ for rs11172113. The *LRP1* gene encodes a precursor protein that is processed by furin in the trans-Golgi complex, generating a 515 kDa alpha-chain and an 85 kDa beta-chain associated non-covalently (25). This protein is involved in multiple cellular processes, including intracellular signalling, lipid homeostasis, and clearance of apoptotic cells (26). The *LRP1* gene has also been reported to be associated with Alzheimer's disease, cardiovascular disease, and tumours (27–29). Chasman et al first identified the association between the *LRP1* gene and migraine in a Caucasian population.^14^ The association was then replicated by Freilinger et al in migraine without aura in German and Dutch patients (15). The locus was further replicated in Indian but not in Chinese populations (30.31). It is worth noting that many nearby supporting SNPs for the *LRP1* region were located in the nearby gene of *STAT6* (which was also strongly associated with headache).

The second SNP cluster was in the *FHL5* gene area with a lowest *P* value of 2.58 × 10^−31^ for rs9486715. The protein encoded by this gene is expressed with activator of cAMP-responsive element modulator (CREM) (32). It is associated with CREM and confers a powerful transcriptional activation function (32). The locus was first reported by Anttila et al to be associated with migraine (13). Although replication failed in a Chinese population, the locus was replicated by Gormley et al in a large meta-analysis on migraine (17,33). Both the *LRP1* and *FHL5* genes are also candidate genes for cervical artery dissection suggesting vascular involvement in headache (18). Just as the *LRP1* region extends to the *STAT6* gene, the *FHL5* cluster extends to the *UFL1* gene.

The strongest association among the 14 newly proposed loci is located in the *LINC02210-CRHR1* gene. The corticotropin releasing hormone receptor 1 (*CRHR1*) gene has been reported to be associated with stress, panic disorder, neuroticism (34–36). This is matched with our genetic correlation results by LD hub suggesting headache and psychological traits share genetic architectures (see discussion below). The loci extends to the microtubule associated protein tau (*MAPT*) gene region. Other new identified loci included: calcium/calmodulin dependent protein kinase ID (*CAMK1D), NUFIP2, FMR1 interacting protein 2 (NUFIP2), butyrophilin subfamily 2 member A2 (BTN2A2), myosin IH (MYO1H), intraflagellar transport 81 (IFT81), nucleolar protein 4 like (NOL4L), polypyrimidine tract binding protein 2 (PTBP2), frataxin (FXN), autophagy related 13 (ATG13), microtubule-actin crosslinking factor 1 (MACF1)* and some intergenic regions. It is interesting to notice that GWAS have identified that *CAMK1D* and *MACF1* are involved in vascular disorders (hypertension and peripheral artery disease) (37,38), again supporting a vascular contribution to headache.

It is also interesting to note that the *STAT6* gene was the most significant in the gene analysis with headache and not the *LRP1* gene where the top SNP resides. These two genes are next to each other in the genome and have previously been associated with disorders related to the immune system such as food allergen sensitization and Sjogren's Disease (39,40). It is reported that IL4/STAT6 signalling activates neural stem cell proliferation and neurogenesis in zebrafish brain, which indicates the importance of the gene in neuron function (41). The gene sets analysis revealed that genes involved in the neurogenesis are associated with headache which is consistent with the tissue expression analysis.

Both tissue expression analysis on 30 general tissue types and 53 specific tissue types showed significant associations between brain tissues and headache, but not vascular tissues. This conclusion is different from the predominant theory of vascular aetiology for migraine since our results suggest that brain function is closely related to broadly-defined headache. Combining all results from GWAS and FUMA together, it is clear that both neuronal and vascular factors are involved in the headache mechanism.

Consistent with previous studies from our group, we found strong evidence to support a shared genetic aetiology of pain with psychological traits, indicating a vulnerability to depression and other negative mood states (42,43). In addition, we found a negative correlation with factors associated with a longer duration of education. Previous results have suggested that headache sufferers were more likely to have psychiatric disorders than healthy people (44). In addition, a shared genetic link between migraine and depression has been identified (45). Further studies are needed to demonstrate the nature of these correlations and whether any directional ‘causal’ inferences can be drawn.

The Q-Q plot suggested that there are residual confounding factors between headache cases and controls which have not been adjusted for. Those could be factors associated with psychiatric and vascular disorders. We also noted the is a 90 degree upswing (around the *P* = 10^−14^ level). This is due to the fact that we have a cluster of 1,800 significant SNPs at this level (mainly located at the *LINC02210-CRHR1-MAPT* loci), which is over half of the total number of significant SNPs.

Using the CaTS power calculator, we had 80% power to identify SNP associations with a significance level of 5 × 10^−8^, based on 74,461 cases and 149,312 controls, assuming an additive model, a minor disease allele frequency of 0.15, a genotypic relative risk of 1.05, and a prevalence of headache in the general population of 0.2 (a conservative assumption) (46).

In conclusion, we have identified 28 loci for broadly-defined headache as a single entity in a GWAS using the UK Biobank resource, including 14 loci that have previously been associated with migraine, and 14 loci that have not previously been associated with headache. This is the largest GWAS on headache in a single population so far. We also identified evidence that brain function is closely related to broadly-defined headache. In addition, we found several significant correlations with a number of psychological factors, suggesting that the genetic aetiology of headache may also be related to these traits.

## Materials and Methods

### Participants and genetic information of participants

The UK Biobank is a health research resource that aims to improve the prevention, diagnosis and treatment of a wide range of illnesses. The UK Biobank cohort recruited over 500,000 people aged between 40-69 years in 2006-2010 across the UK. Participants completed a detailed clinical, demographic, and lifestyle questionnaire, underwent clinical measures, provided biological samples (blood, urine and saliva) for future analysis, and agreed to have their health records accessed. The informed consent of all participants has been obtained. Details of the UK Biobank resource can be found at www.ukbiobank.ac.uk.

UK Biobank received ethical approval from the National Health Service National Research Ethics Service (reference 11/NW/0382). The current analyses were conducted under approved UK Biobank data application number 4844.

The detailed methods of DNA extraction and quality control can be found at http://www.ukbiobank.ac.uk/wp-content/uploads/2014/04/DNA-Extraction-at-UK-Biobank-October-2014.pdf. Participants’ DNA was genotyped by bespoke Affymetrix UK Biobank chips. Standard quality-control (QC) steps were performed by the Wellcome Trust Centre for Human Genetics at Oxford University. In this study, we used BGENIE (https://jmarchini.org/bgenie/) to be the main GWAS software and removed SNPs with INFO scores less than 0.1, with minor allele frequency less than 0.5%, or those that failed Hardy-Weinberg tests *P* < 10^−6^ (47). SNPs on the X and Y chromosomes and mitochondrial SNPs were excluded from analyses. The detailed QC steps can be found at http://biobank.ctsu.ox.ac.uk/crystal/refer.cgi?id=155580.

In September 2017, the genetic information from 501,708 samples was released to UK Biobank project research collaborators. This included imputation with a reference set combining the UK10K haplotype and 1000 Genomes Phase 3 reference panels. The imputation details can be found at http://biobank.ctsu.ox.ac.uk/crystal/refer.cgi?id=157020.

### Phenotypic information on pain

The UK Biobank participants were offered a pain-related questionnaire, which included the question: ‘in the last month have you experienced any of the following that interfered with your usual activities?’. The options were: 1. Headache; 2. Facial pain; 3. Neck or shoulder pain; 4. Back pain; 5. Stomach or abdominal pain; 6. Hip pain; 7. Knee pain; 8. Pain all over the body; 9. None of the above; 10. Prefer not to say. Participants could select more than one option. (UK Biobank Questionnaire field ID: 6159)

The headache cases in this study were those who selected the ‘Headache’ option for the above question, regardless of whether they had selected other options.

The controls in this study were those who selected the ‘None of the above’ option.

### Statistical analysis

Standard Frequentist association tests using BGENIE was used to perform association studies adjusting for age, sex, BMI, 9 population principal components, genotyping arrays, and assessment centres. Gender difference between cases and controls was compared using chi-square testing. Age and BMI were compared using independent t testing in IBM SPSS 22 (IBM Corporation, New York). SNP associations were considered significant if they had a *P* value less than 5 × 10^−8^.

SNP functional annotations were applied by the FUMA web application and a Manhattan plot was generated by R (48). R was also used to generate the corresponding Q-Q plot, a tool to evaluate differences between cases and controls caused by potential confounders.

The gene analysis and gene-set analysis were performed with MAGMA v1.6, which was integrated in FUMA (49). Both analyses were based on GWAS summary statistics. In gene analysis, summary statistics of SNPs are aggregated to the level of whole genes, testing the joint association of all SNPs in the gene with the phenotype. In gene-set analysis, individual genes are aggregated to groups of genes sharing certain biological, functional or other characteristics. This will provide insight into the involvement of specific biological pathways or cellular functions in the genetic aetiology of a phenotype. Tissue expression analysis was obtained from GTEx (https://www.gtexportal.org/home/) which was also integrated in FUMA.

In order to identify genetic correlations with other complex traits and diseases, we used linkage disequilibrium score regression through LD Hub v1.4.1 (available at http://ldsc.broadinstitute.org/ldhub/) (50). This web-tool uses individual SNP allele effect sizes and the average linkage disequilibrium in a region to estimate the bivariate genetic correlations of headache with 234 traits. Those with *P* values of 2.1 × 10^−4^ (0.05/234) or less should be regarded as surviving Bonferroni adjustment for multiple testing.

## Acknowledgements/Funding

This work was supported by the DOLORisk project [EU Horizon2020, grant number: 633491], the STRADL project [Wellcome Trust, grant number: 104036/Z/14/Z], and the Centre for Cognitive Ageing and Cognitive Epidemiology [Medical Research Council and Biotechnology and Biological Sciences Research Council, grant number: MR/K026992/1]. We are grateful for support from the the Dr Mortimer And Theresa Sackler Foundation.

We would like to thank all participants of the UK Biobank cohort who have provided necessary genetic and phenotypic information.

## Supporting information

Supplementary Fig. S1: The Q-Q plot of the GWAS on headache using UK Biobank Supplementary Table S1: A summary of 3343 GWAS significant SNPs

Supplementary Table S2: Significant genes based on the gene analysis by MAGMA integrated in FUMA

Supplementary Table S3: Top 10 gene sets results by MAGMA integrated in FUMA

Supplementary Table S4: Tissue expression analysis on 30 general tissue types

Supplementary Table S5: Tissue expression analysis on 53 specific tissue types

Supplementary Table S6: Genetic correlations between headache and other phenotypes which reached significance of 0.05/234

